# Locomotor and Reinforcing Effects of Pentedrone, Pentylone and Methylone in Rats

**DOI:** 10.1101/166579

**Authors:** Mehrak Javadi-Paydar, Jacques D. Nguyen, Sophia A. Vandewater, Tobin J. Dickerson, Michael A. Taffe

**Affiliations:** Department of Neuroscience; Department of Chemistry; The Scripps Research Institute; La Jolla, CA, USA

**Author notes:** Address Correspondence to: Dr. Michael A. Taffe, Department of Neuroscience, SP30-2400; 10550 North Torrey Pines Road; The Scripps Research Institute, La Jolla, CA 92037; USA.

## Abstract

The broad diversity of synthetic cathinone psychostimulant drugs that are available to users complicates research efforts to provide understanding of health risks. Second generation cathinones pentedrone and pentylone are distinguished from each other by the 3,4-methylenedioxy structural motif (which distinguishes methamphetamine from 3,4-methylenedioxymethamphetamine) and each incorporates the α-alkyl chain motif contained in the transporter-inhibitor cathinones (3,4-methylenedioxypyrovalerone (MDPV), α-pyrrolidinopentiophenone (α-PVP)) but not in the monoamine releasers (mephedrone, methylone). Studies were conducted in male and female Wistar rats to compare locomotor and thermoregulatory effects of pentedrone, pentylone and methylone using an implanted radiotelemetry system. Reinforcing effects were assessed in female Wistar rats trained in the intravenous self-administration (IVSA) procedure and subjected to dose-substitution (0.025-0.3 m/gkg/inf) under a fixed-ratio 1 response contingency. Pentedrone, pentylone and methylone dose-effect curves were contrasted with those for α-PVP and α-pyrrolidinohexiophenone (α-PHP). Dose dependent increases in locomotion were observed after intraperitoneal injection of pentylone (0.5-10.0 mg/kg), pentedrone (0.5-10.0 mg/kg) or mephedrone (0.5-10.0 mg/kg) in male and female rats. The maximum locomotor effect was similar across drugs but lasted longest after pentedrone. Mean body temperature did not vary systematically more than 0.5 °C after pentedrone or pentylone in either sex. A sustained hyperthermia (0.4-0.8 °C) was observed for four hours after 10 mg/kg methylone in male rats. More infusions of pentedrone or pentylone were self-administered compared with methylone, but all three were less potent than α-PVP or α-PHP. These studies support the inference that second generation cathinones pentylone and pentedrone have abuse liability greater than that of methylone.

## 1. Introduction

Recreational use of cathinone derivative psychostimulant drugs has increased over the past decade (Madras, 2017), yet the broad diversity of the drugs that are available to users (Brunt *et al*, 2017; Odoardi *et al*, 2016) has complicated research efforts to provide understanding of various health risks (Aarde and Taffe, 2017; Angoa-Perez *et al*, 2017; Negus and Banks, 2017; Papaseit *et al*, 2017). The US DEA placed both pentedrone and pentylone under temporary Schedule I control in March of 2014 and this action was finalized in 2017 (Drug Enforcement Administration, 2014, 2017). These compounds have been detected in forensic casework (Adamowicz *et al*, 2016; Elliott and Evans, 2014), and there is evidence for health threatening toxic effects of pentylone (Liakoni *et al*, 2015) as well as for pentedrone in combination with other cathinone derivatives (Liveri *et al*, 2016; Sykutera *et al*, 2015). Second-generation compounds such as pentylone and pentedrone have received less research attention compared with mephedrone, methylone, 3,4-methylenedioxypyrovalerone (MDPV) and α-pyrrolidinopentiophenone (α-PVP).

Although prior preclinical evidence is not comprehensive, there is evidence that pentedrone increases locomotor activity in mice (Gatch *et al*, 2015; Hwang *et al*, 2017), supports intravenous self-administration (IVSA) in male Wistar rats (Hwang *et al*, 2017) and substitutes for the discriminative stimulus effect of methamphetamine and cocaine in male rats (Gatch *et al*, 2015). Pentylone likewise increases locomotor activity in mice and substitutes for methamphetamine and cocaine (Gatch *et al*, 2015). In contrast the effects of methylone are better established. Methylone supports IVSA in both male and female rats (Aarde *et al*, 2015b; Creehan *et al*, 2015; Nguyen *et al*, 2016b; Schindler *et al*, 2015; Vandewater *et al*, 2015; Watterson *et al*, 2012), facilitates intracranial self-stimulation reward in rats (Bonano *et al*, 2014), can condition a place preference in mice (Karlsson *et al*, 2014), substitutes for the discriminative stimulus effects of cocaine or methamphetamine in rats (Gatch *et al*, 2013) and increases body temperature and locomotor activity in rats and mice (Gatch *et al*, 2013; Grecco and Sprague, 2016; Kiyatkin *et al*, 2015; Marusich *et al*, 2012). Despite the fact that methylone is structurally similar to 3,4-methylenedioxymethamphetamine (MDMA) the pre-clinical self-administration data suggests that methylone exhibits enhanced liability for compulsive use compared with that of MDMA (Nguyen *et al*, 2016b; Vandewater *et al*, 2015; Watterson *et al*, 2012).

The diversity of cathinone derivatives permits further investigation of the role of various substitution moieties common to both amphetamine and cathinone drugs of abuse. In this study, we investigate the 3,4-methylenedioxy motif (see **Figure 1**) in the contrast of the effects of pentedrone with pentylone. This motif, when added to methamphetamine to produce MDMA, confers reduced rewarding potency and efficacy (Dalley *et al*, 2007; Schenk, 2009; Vandewater *et al*, 2015), reduced locomotor potency (Huang *et al*, 2012; Miller *et al*, 2013a), reduced efficacy to induce stereotyped, repetitive behavior and increased thermoregulatory disruption (Miller *et al*, 2013a). In contrast, the presence of the 3,4-methylenedioxy substitution produces no change *in vivo* in the context of the closely related, restricted transporter inhibitor cathinones α-PVP and MDPV which exhibit similar efficacy and potency on both locomotor and self-administration assays in rats (Aarde *et al*, 2015a). Pentedrone and pentylone also include the extended alkyl-tail carbon chain that is present on MDPV and α-PVP which may be related to the restriction of those drugs to transporter inhibition. This might predict that the 3,4-methylenedioxy motif has minimal impact on these additional compounds which lack the pyrrolidine ring of MDPV and α-PVP.

**Figure 1:**
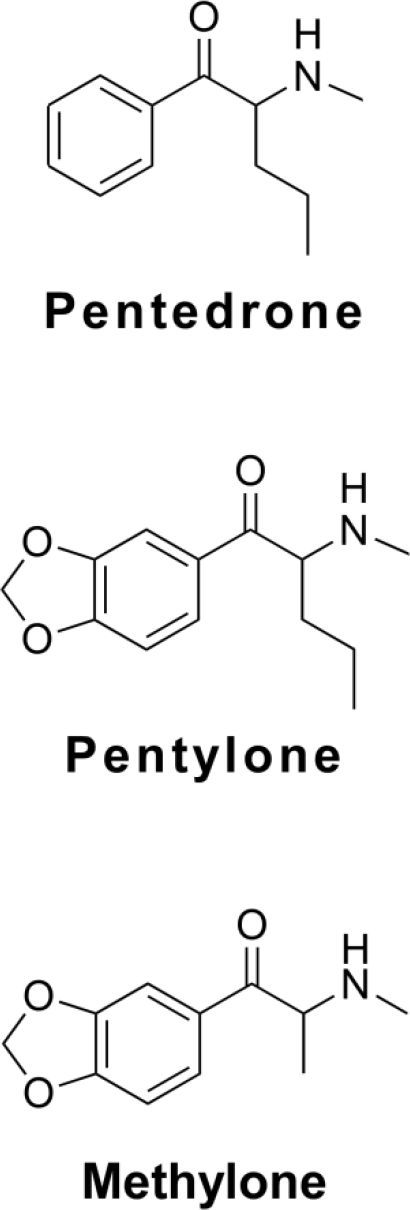
Chemical structures of the substituted cathinones under investigation.

Pentedrone and pentylone exhibit negligible dopamine or norepinephrine release (Eshleman *et al*, 2017; Simmler *et al*, 2014), however some serotonin release was observed for pentylone in one of the reports and methylone is more effective than either compound (Simmler *et al*, 2014). The inhibition of the serotonin transporter (SERT) by pentylone is 16-fold higher than inhibition by pentedrone, similar to the 17-fold higher affinity of MDMA over methamphetamine. In addition, pentylone inhibits dopamine transporter (DAT) activity with a potency of about half that of pentedrone whereas DAT inhibition caused by methamphetamine is 17-fold higher than that of MDMA. The ratio of potencies for inhibiting the DAT versus the SERT is 54 for pentedrone, compared with 6.2 for pentylone, 3.2 for methylone, 22 for methamphetamine and 0.08 for MDMA (Simmler *et al*, 2013; Simmler *et al*, 2014). The *in vitro* pharmacology therefore predicts that pentedrone would be significantly more potent as a locomotor stimulant or reinforcer compared with pentylone and methylone; however, the difference should be less pronounced than the potency difference between MDMA and methamphetamine. The potential of pentylone to release serotonin predicts reduced efficacy compared with pentedrone. The present study was conducted to evaluate these hypotheses *in vivo* using IVSA techniques (Aarde *et al*, 2013a; Aarde *et al*, 2015a; Creehan *et al*, 2015) and a rat locomotor assay that has been used to evaluate the locomotor stimulant effects of MDPV, α-PVP, methamphetamine and MDMA (Aarde *et al*, 2015a; Miller *et al*, 2013a). In these procedures, comparison of behavioral outcomes after a range of doses can be used to infer differences in potency (the dose at which a given effect occurs) and efficacy (the maximum effect observed for any dose). These differences, if established, can then be used to support predictions or inferences about the relative abuse liability when translated to the recreational use context.

## 2. Methods

### 2.1 Subjects

Male (N=8) and female (N=33) Wistar (Charles River, New York) rats entered the laboratory at 10 weeks of age and were housed in humidity and temperature-controlled (23±1 °C) vivaria on 12:12 hour light:dark cycles. Experimental procedures took place during the animals’ dark cycle. Animals had ad libitum access to food and water in their home cages. All procedures were conducted under protocols approved by the Institutional Care and Use Committees of The Scripps Research Institute and in a manner consistent with the Guide for the Care and Use of Laboratory Animals (National Research Council (U.S.). Committee for the Update of the Guide for the Care and Use of Laboratory Animals. *et al*, 2011).

### 2.2 Drugs

Drugs were dissolved in physiological saline for the i.p. or i.v. routes of administration. Pentylone HCl, pentedrone HCl, α-pyrrolidinopentiophenone HCl (α-PVP) and α-pyrrolidinohexiophenone HCl (α-PHP) were obtained from Cayman Chemical. Methylone HCl was obtained from NIDA Drug Supply. Dosing is expressed as the salt.

### 2.3 Procedures

#### 2.3.1 Surgery

##### 2.3.1.1 Radiotransmitter Implantation

Rats were anesthetized with an isoflurane/oxygen vapor mixture (isoflurane 5% induction, 1-3% maintenance), and sterile radiotelemetry transmitters (Data Sciences International; TA-F40) were implanted in the abdominal cavity through an incision along the abdominal midline posterior to the xyphoid space as previously reported (Aarde *et al*, 2015a; Miller *et al*, 2013a; Wright *et al*, 2012). Absorbable sutures were used to close the abdominal muscle incision and the skin incision was closed with the tissue adhesive (3M Vetbond Tissue Adhesive; 3M, St Paul, MN). A minimum of 7 days was allowed for surgical recovery prior to starting experiments. For the first three days of the recovery period, an antibiotic Cefazolin (Hikma Farmaceutica, Portugal; 0.4 mg/kg, i.m. in sterile water Day 1, s.c. Day 2-3) and an analgesic flunixin (FlunixiJect, Bimeda USA, Oakbrook Terrace, IL; 2.5 mg/kg, s.c. in saline) were administered daily.

##### 2.3.1.2 Intravenous catheter implantation

Rats were anesthetized with an isoflurane/oxygen vapor mixture (isoflurane 5 % induction, 1-3 % maintenance) and prepared with chronic intravenous catheters as described previously (Aarde *et al*, 2015a; Aarde *et al*, 2013b; Miller *et al*, 2013b; Nguyen *et al*, 2016b). Briefly, the catheters consisted of a 14-cm length polyurethane based tubing (MicroRenathane®, Braintree Scientific, Inc, Braintree MA, USA) fitted to a guide cannula (Plastics one, Roanoke, VA) curved at an angle and encased in dental cement anchored to an ~3-cm circle of durable mesh. Catheter tubing was passed subcutaneously from the animal's back to the right jugular vein. Catheter tubing was inserted into the vein and secured gently with suture thread. A liquid tissue adhesive was used to close the incisions (3M™ Vetbond™ Tissue Adhesive; 1469S B). A minimum of 4 days was allowed for surgical recovery prior to starting an experiment. For the first 3 days of the recovery period, an antibiotic (cephazolin) and an analgesic (flunixin) were administered daily. During testing and training, intra-venous catheters were flushed with ~0.2–0.3 ml heparinized (32.3 USP/ml) saline before sessions and ~0.2–0.3 ml heparinized saline containing cefazolin (100 mg/ml) after sessions. Starting in the second week of IVSA training, the catheter patency was assessed after the last session of the week via administration through the catheter of ~0.2 ml (10 mg/ml) of the ultra-short-acting barbiturate anesthetic Brevital sodium (1 % methohexital sodium; Eli Lilly, Indianapolis, IN). Animals with patent catheters exhibit prominent signs of anesthesia (pronounced loss of muscle tone) within 3 s after infusion. Animals that failed to display these signs were considered to have faulty catheters and were discontinued from the study. Data that were collected prior to failing this test and after the previous passing of this test were excluded from analysis.

#### 2.3.2 Radiotelemetry Measures of Locomotor Activity and Body Temperature

Locomotor activity and temperature data were collected while animals were housed in clean standard plastic home-cages (thin layer of bedding) in a dark testing room (dim red-light illumination), separate from the vivarium, during the (vivarium) dark cycle. Radiotelemetry transmissions were collected via receiver plates (Data Sciences International; RPC-1) placed under the cages as described in prior investigations (Aarde *et al*, 2013b; Miller *et al*, 2013a; Taffe *et al*, 2015; Wright *et al*, 2012). The ambient temperature for the studies was 22 ±1 °C. Sessions started with a 30-minute interval in the recording cage to determine a pre-treatment baseline of activity and temperature. Thereafter animals were injected with the scheduled challenge drug/vehicle and then returned to their individual recording cages. The three telemetry samples taken prior to moving the rat to the inhalation chamber were used as the pre-treatment baseline for analysis. Primary analysis focused on the average activity rate (counts per minute) and body temperature (°C) in 30 minute intervals as derived from the primary 5 min sampling bins. Data are timed to the injection and the “30 minute” time bin reflects the average of 6 samples collected from 5 to 30 minutes after return to the recording chamber following injection. Any missing body temperature data (e.g., due to radio interference or animal's location within the chamber at the time of sampling) was interpolated across preceding and succeeding recorded values.

### 2.4 Experiments

#### 2.4.1 Temperature and Activity in Male Rats

Male Wistar rats (Group 1; N=8; 33 weeks of age and a mean of 589 g at the start of this study) were initially exposed (~15 weeks of age) to inhalation of propylene glycol (PG) vapor versus MDMA (400 mg/ml in PG) for 40 minutes in a method previously described (Nguyen *et al*, 2016a). No other experiments were performed with this group until the current experiment. Rats were tested first with pentedrone (0.0, 0.5, 1.0, 5.0 mg/kg, i.p.), then α-PPP (0.0, 0.5, 1.0, 5.0 mg/kg, i.p.) and then pentylone (0.0, 0.5, 1.0, 5.0 mg/kg, i.p.) with dose order randomized within-drug compound. Doses were initially selected based on prior and ongoing locomotor studies in the laboratory with compounds with a similar range of *in vitro* pharmacological effects. Following this, animals were tested with 10 mg/kg, i.p. of pentedrone, α-PPP, and pentylone in a randomized order to further determine the dose-effect range. Next, rats were tested with saline and 5.0 mg/kg, i.p. of α-PVP, methamphetamine or MDPV in a randomized order. One animal was euthanized after the methamphetamine condition by protocol due to unregulated elevated body temperature, thus N=7 for the subsequent studies. The final experiment was to evaluate the effects of treatment with methylone (0.0, 0.5, 1.0, 5.0 mg/kg, i.p.) in a randomized order followed by methylone (10 mg/kg, i.p.).

#### 2.4.2 Temperature and Activity in Female Rats

Female Wistar rats (Group 2; N=8; 12 weeks of age and a mean of 222 g at the start of this study) were initially recorded in a habituation session with no treatments. For this study, rats were evaluated first with α-PPP (0.0, 1.0, 5.0, 10.0 mg/kg, i.p.), then pentedrone (0.0, 1.0, 5.0, 10.0 mg/kg, i.p.) then pentylone (0.0, 1.0, 5.0, 10.0 mg/kg, i.p.) and finally methylone (0.0, 1.0, 5.0, 10.0 mg/kg, i.p.).

#### 2.4.3 Intravenous Self-Administration in Female Rats

Female Wistar rats (Group 3; N=17; 10 weeks of age at the start of this study) were prepared with intravenous catheters and trained to self-administer α-PVP (N=8; 0.05 mg/kg per infusion) or pentedrone (N=9; 0.2 mg/kg/inf) using a fixed-ratio 1 (FR1) response contingency. Doses were selected from prior self-administration studies and ongoing locomotor studies to result in similar numbers of infusions throughout the acquisition interval. Following 13 sessions of acquisition the training doses were halved for another seven sessions (data not shown). Thereafter, animals were subjected to dose substitution of different cathinones (0.0, 0.025, 0.05, 0.1, 0.3 mg/kg/inf) in a randomized order, with the restricted transporter-inhibitor compounds serving as positive controls / validation studies. The animals first completed a dose-substitution series with their respective training drug. The pentedrone trained animals next completed the series in the following order: α-PPP, α-PVP, pentylone, α-PHP, methylone. The α-PVP group next completed additional series as follows: α-PPP, pentedrone, pentylone, α-PHP, methylone. A follow-up study including saline and doses (0.0125, 0.025, 0.1 mg/kg/inf) of α-PVP and α-PHP was conducted to further determine the dose-effect range for these two compounds.

### 2.5 Data Analysis

Measures of locomotor activity (rate of activity counts per minute) were analyzed by repeated-measures Analysis of Variance (rmANOVA) with Time Post-injection and Drug Dose (or Drug for the 10 mg/kg comparison) as within-subjects factors. Body temperature data were not formally analyzed as mean changes of >5°C were only observed sporadically. The number of infusions obtained in the IVSA experiments was analyzed by rmANOVA with Sessions (acquisition only), Drug identity and/or Dose (dose-substitution only) as within-subjects factors. Significant main effects from the rmANOVA were further analyzed with post hoc multiple comparisons analysis using the Tukey procedure. The criterion for significant results was at P < 0.05 and all analyses were conducted using Prism 6 or 7 for Windows (v. 6.02, v. 7.00; GraphPad Software, Inc, San Diego CA).

## 3. Results

### 3.1 Effect of Substituted Cathinones on Activity and Body Temperature in Male Rats

#### 3.1.1 Activity

Intraperitoneal injection of pentylone, pentedrone and methylone all significantly increased locomotor activity in a dose-dependent manner in the male rats (**Figure 2**).

**Figure 2:**
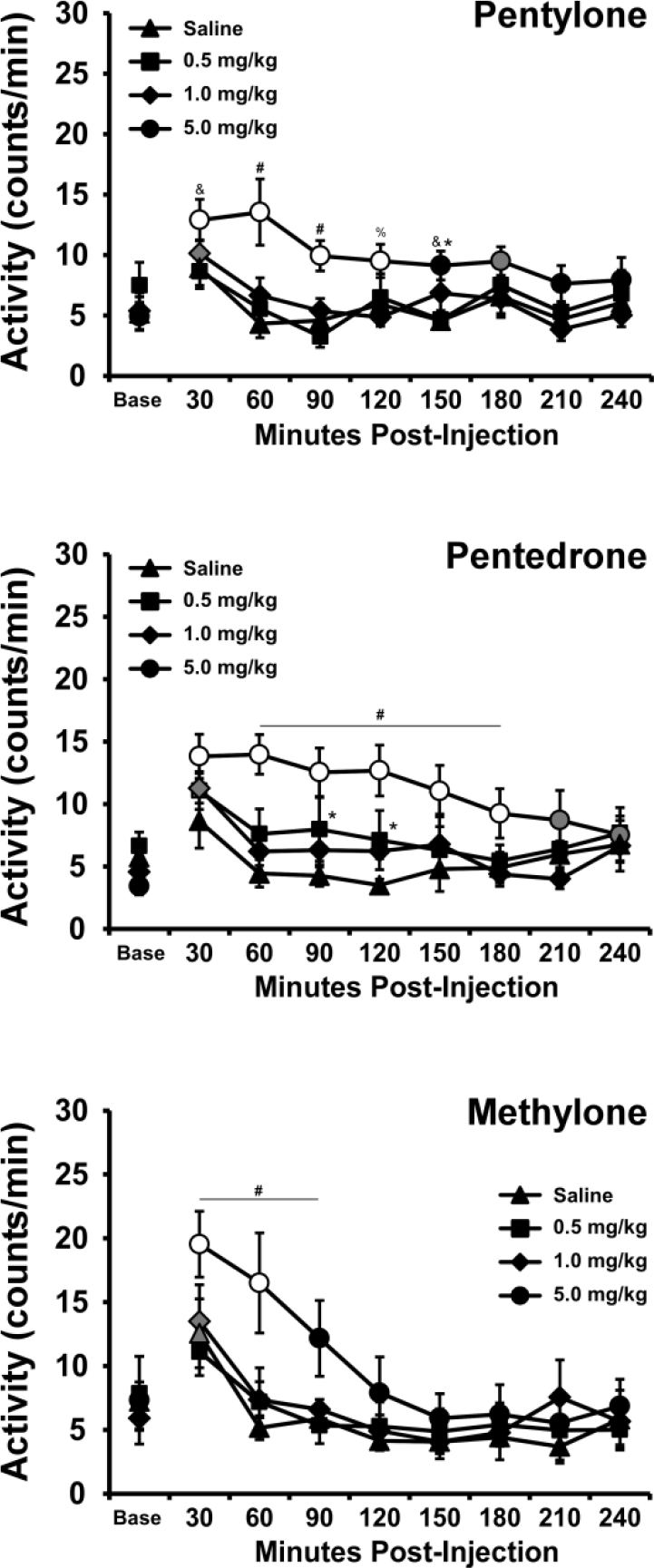
Mean (±SEM) activity rates of male rats following i.p. injection with pentylone (N=8), pentedrone (N=8) and methylone (N=7). Significant differences from the baseline, within drug/dose, are indicated by symbols filled with gray and from both vehicle and baseline with open symbols. Significant differences from 0.5 mg/kg and 1.0 mg/kg doses are indicated by #, from 0.5 mg/kg by &, from 1.0 mg/kg by % and from vehicle (only) by *. Base = baseline.

##### 3.1.1.1 Pentylone

The analysis confirmed a significant effect of Time [F (8, 56) = 11.71; P<0.0001], of Dose [F (3, 21) = 12.63; P<0.0001] and of the interaction of factors [F (24, 168) = 2.39; P=0.0007]. The post-hoc test further confirmed that activity was significantly higher after injection of 5.0 mg/kg pentylone compared with the vehicle (30-150 minutes post-injection), with the 0.5 mg/kg dose (30-90, 150 minutes post-injection) and with the 1.0 mg/kg dose (60-120, 210 minutes post-injection).

##### 3.1.1.2 Pentedrone

The analysis confirmed a significant effect of Time [F (8, 56) = 3.43; P<0.005]. The post-hoc test confirmed activity was significantly higher relative to baseline 30-90 minutes after injection of 5.0 mg/kg pentedrone but not for any other of the conditions.

##### 3.1.1.3 Methylone

The analysis confirmed a significant effect of Time [F (8, 48) = 26.26; P<0.0001] and of the interaction of Time with Dose [F (24, 144) = 2.66; P<0.0005]. The post-hoc test further confirmed that activity was significantly higher 30-90 minutes after injection of 5.0 mg/kg Methylone compared with all other conditions.

##### 3.1.1.4 Pentylone/Pentedrone/Methylone (10 mg/kg)

The analysis confirmed significant effects of Time [F (8, 216) = 32.0; P<0.00011], of Drug [F (3, 27) = 5.99; P<0.005] and of the interaction of Time with Drug [F (24, 216) = 6.63; P<0.0001], see **Figure 3**. The post-hoc test furthermore confirmed that activity was significantly elevated over the vehicle condition after 10 mg/kg pentylone [30-90 minutes post-injection], after 10 mg/kg pentedrone [30-180 minutes post-injection] and after 10 mg/kg methylone (30-120 minutes post-injection). Furthermore, activity was significantly higher after methylone compared with pentylone 90 minutes post-injection.

**Figure 3:**
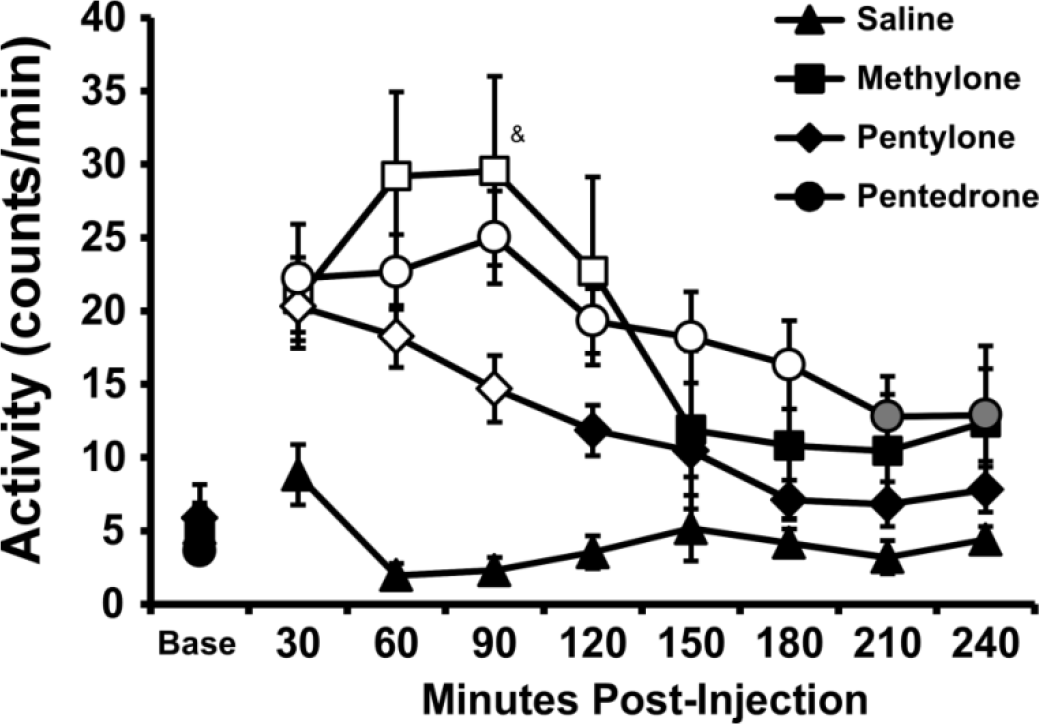
Mean (±SEM) activity rates of male rats following injection with the vehicle or 10 mg/kg, i.p. pentylone (N=8), pentedrone (N=8) and methylone (N=7). Sig differences from the baseline, within drug/dose, are indicated by symbols filled with gray and from both vehicle and baseline with open symbols. A significant difference from pentylone is indicated by &. Base = baseline.

#### 3.1.2 Temperature

Body temperature was not substantially changed in most of these studies; in the vast majority of cases mean body temperature did not vary more than 0.5°C from the baseline temperature (data not shown). Episodic temperature change in excess of a 0.5°C deviation included mean temperature changes of +0.58°C at 60 min after 5.0 mg/kg pentedrone, +0.51°C 60 minutes after 5.0 mg/kg pentylone and −0.52°C 90 minutes after 0.5 mg/kg pentylone. Sustained change was only observed after the 10 mg/kg methylone dose, following which mean elevations of 0.71-0.84°C were observed 60-120 minutes after injection and of 0.53-0.57°C from 150-240 minutes after injection.

### 3.2 Effect of Substituted Cathinones on Activity and Body Temperature in Female Rats

Intraperitoneal injection of pentylone, pentedrone and methylone all significantly increased locomotor activity in a dose-dependent manner in the female rats (**Figure 4**).

**Figure 4:**
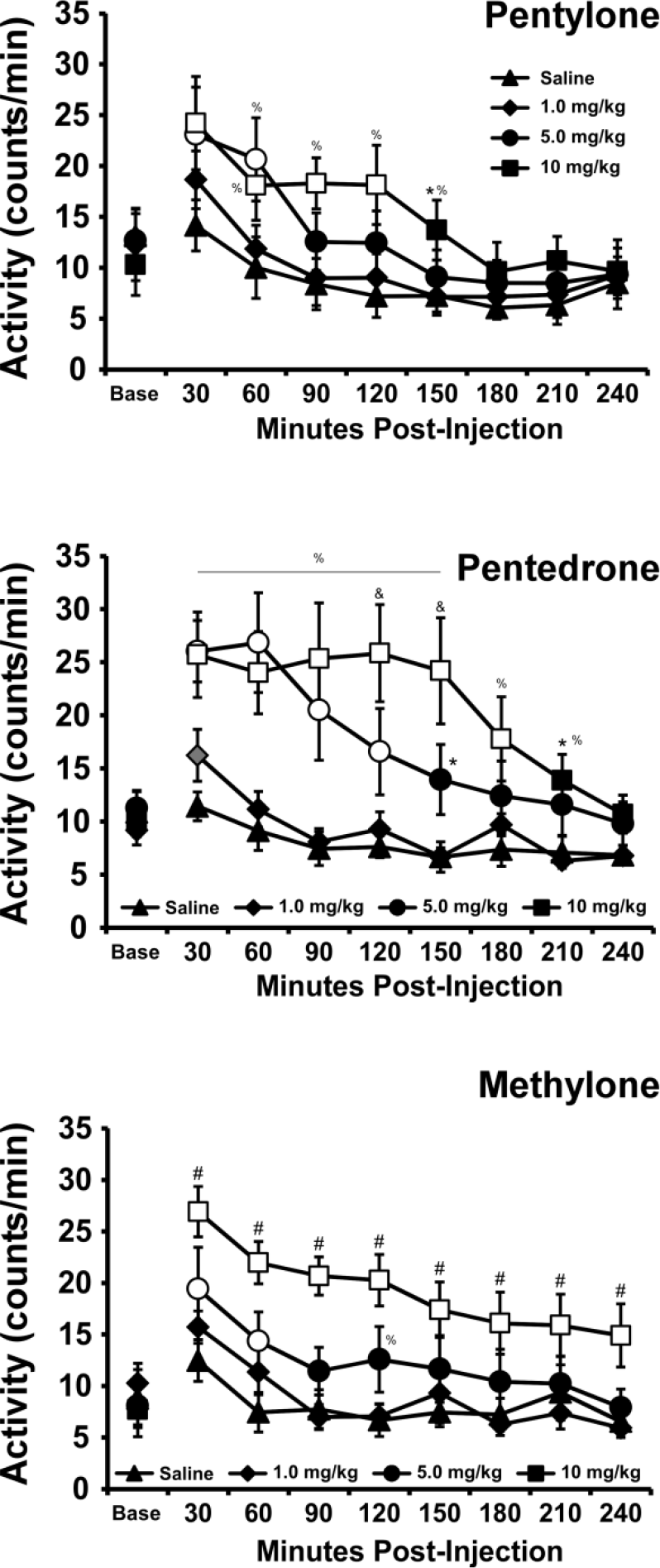
Mean (±SEM) activity rates of female rats (N=8) following injection with pentylone, pentedrone or methylone. Significant differences from the baseline, within drug/dose, are indicated by symbols filled with gray and differences from both vehicle and baseline with open symbols. Significant differences from all other dose conditions are indicated with #, from the 5 mg/kg dose with & and from 1 mg/kg by % (line indicates both 5 and 10 mg/kg differ from 1 mg/kg). Base = baseline.

#### 3.2.1 Activity

##### 3.2.1.1 Pentylone

The analysis confirmed significant effects of time [F (8, 56) = 16.69; P<0.0001], of dose [F (3, 21) = 7.91; P<0.005] and an interaction of time with dose [F (24, 168) = 2.24; P<0.005] on activity after injection of pentylone (**Figure 4**). The post-hoc test further confirmed that activity was significantly higher compared with vehicle after 5.0 mg/kg dose (30-60 minutes after injection) or the 10.0 mg/kg dose (30-150 minutes), and higher than 1 mg/kg dose after the 5 mg/kg (60 minutes) or 10 (60-150) mg/kg doses.

##### 3.2.1.2 Pentedrone

Activity rates after injection of pentedrone (**Figure 4**) were significantly affected by time [F (8, 56) = 24.25; P<0.0001], by dose [F (3, 21) = 12.08; P<0.0001] and the interaction of time with dose [F (24, 168) = 5.86; P<0.0001]. The post-hoc test confirmed activity was significantly higher after 5.0 or 10.0 mg/kg relative to vehicle or 1.0 mg/kg, i.p. from 30-150 minutes after injection. The activity increase after the 10 mg/kg dose remained significantly different from the vehicle or 1.0 mg/kg conditions to 210 minutes post-injection and was higher compared with the 5.0 mg/kg dose from 120-150 minutes post-injection.

##### 3.2.1.3 Methylone

There was a significant effect of time [F (8, 56) = 27.5; P<0.0001], of dose [F (3, 21) = 28.23; P<0.0001] and an interaction of time with dose [F (24, 168) = 3.23; P<0.0001] on activity following methylone injection (**Figure 4**). The post-hoc test confirmed activity was significantly higher compared with vehicle after 5.0 mg/kg dose (30-60 minutes after injection) and higher than after all three other dosing conditions following the 10.0 mg/kg dose (30-240 minutes). Activity was also significantly higher 120 minutes after the 5 mg/kg dose compared with the 1 mg/kg dose.

#### 3.2.2 Temperature

Body temperature was not substantially changed in the female rats for any of these studies (data not shown). Mean body temperature did not vary more than 0.5°C from the baseline temperature in any of the dosing conditions.

### 3.3 Intravenous Self-Administration of Substituted Cathinones in Female Rats

#### 3.3.1 Acquisition

The two groups of female rats self-administered similar numbers of infusions, i.e. of pentedrone (0.2 mg/kg) and α-PVP (0.05), during acquisition (**Figure 5**). The ANOVA confirmed a significant effect of Sessions of acquisition [F (12, 180) = 9.06; P<0.0001], but not of Drug or of the interaction of factors.

**Figure 5:**
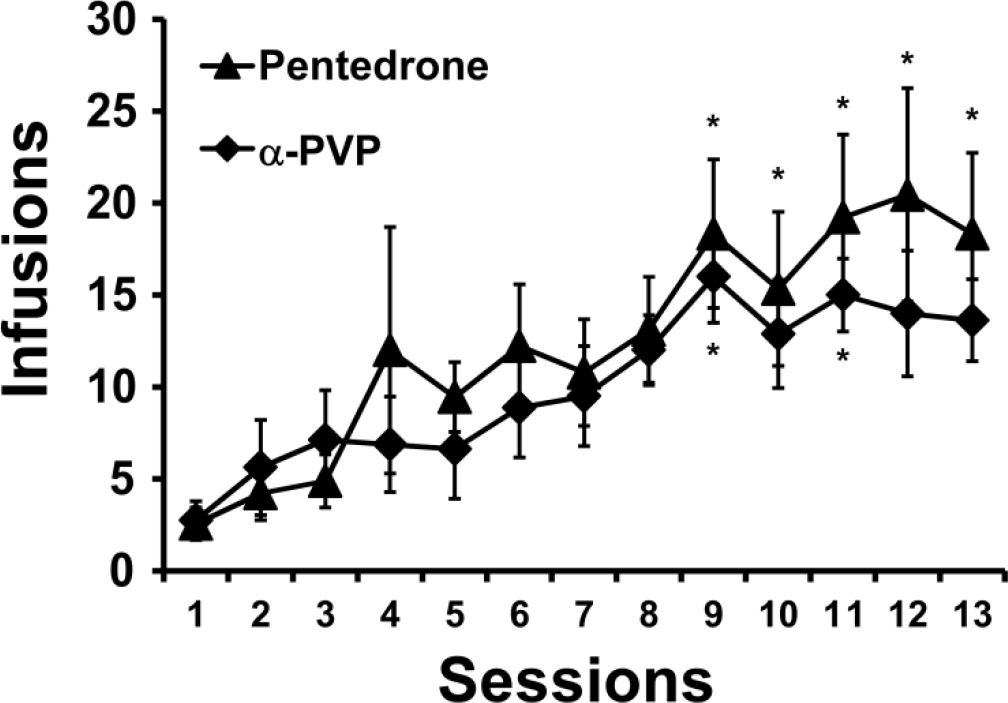
Mean (±SEM) infusions of of pentedrone (0.2 mg/kg) (N=9) and α-PVP (0.05) (N=8) obtained during the initial acquisition sessions. A significant difference from the first session indicated by *.

#### 3.3.2 Dose-Substitution

A total of N=10 (N=6 pentedrone-trained, N=4 α-PVP-trained) rats retained patent catheters throughout all of the initial dose-substitution studies. Ascending and descending limbs of the dose-effect function were observed for pentedrone, pentylone and methylone, whereas a monotonic descending limb was observed for α-PVP and α-PHP (**Figure 6**). Statistical analysis of the pentedrone, pentylone and methylone data confirmed significant effects of Drug [F (2, 18) = 4.42; P<0.05] and of Dose [F (4, 36) = 12.35; P<0.0001] on infusions obtained. The post-hoc test confirmed that significantly more infusions were obtained for at least one dose of each drug compared with vehicle and significantly more infusions of pentedrone and pentylone compared with methylone for at least one dose of each drug (see **Figure 6**). Statistical analysis of the α-PVP and α-PHP dose series confirmed significant effects of Drug [F (1, 9) = 25.59; P<0.001], of Dose [F (4, 36) = 16.41; P<0.0001] and of the interaction of Drug with Dose [F (4, 36) = 3.47; P<0.05] on infusions obtained. The post-hoc test confirmed that increased infusions of α-PHP were obtained, compared with α-PVP, at the 0.025 and 0.05 mg/kg/inf doses.

**Figure 6:**
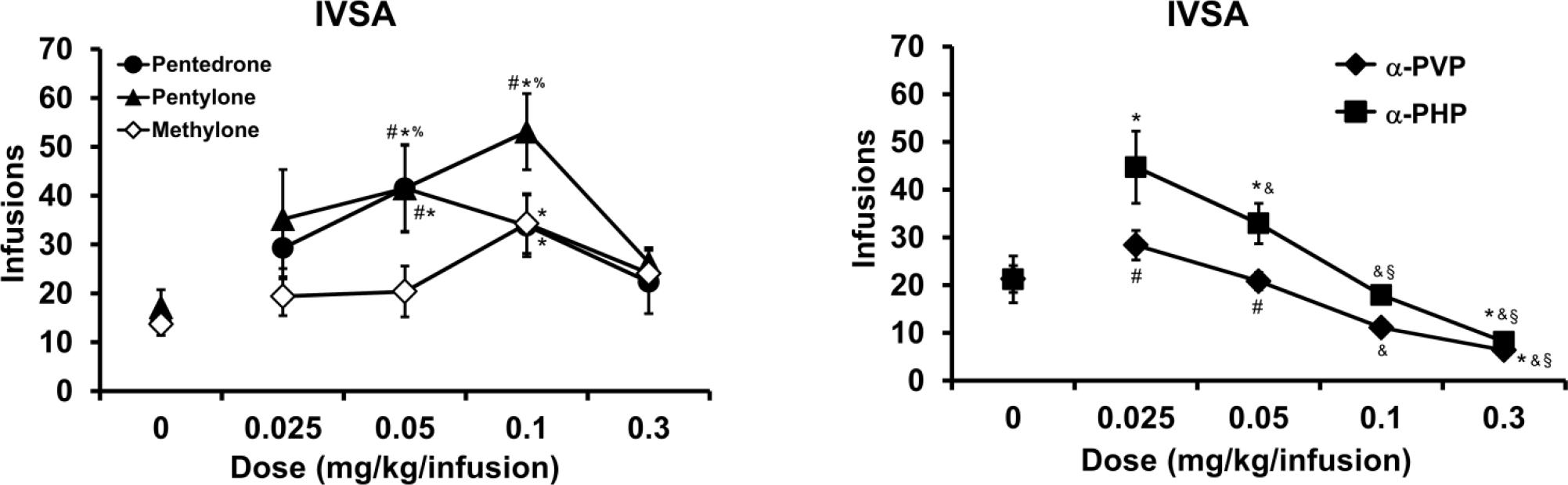
Mean (N=10; ±SEM) infusions of Left) pentylone, pentedrone and methylone and Right) α-PVP and α-PHP obtained by female rats. A significant difference from vehicle is indicated with *, from the 0.025 mg/kg dose with &, from the 0.05 mg/kg dose with §, from the 0.3 mg/kg dose by % and a significant difference from methylone and α-PHP, respectively, is indicated with #.

A follow-up dose-substitution was conducted to evaluate lower dose (0.0125) of α-PVP and α-PHP (Figure 7); a total of N=5 completed this final phase of the study with patent catheters. The primary analysis compared infusions obtained in seven conditions for this final dose series (i.e., saline and 0.0125, 0.025, 0.1 mg/kg/inf for each of the two compounds). The ANOVA confirmed a main effect of dose [F (3, 12) = 26.47; P<0.0001] but not of drug identity or any interaction; the Tukey post-hoc effects are summarized on Figure 7. Because only part of the original group completed the follow-up, the original curves for this subset of animals is graphed for comparison. Statistical analysis within-drug did not confirm any difference in infusions obtained for the overlapping doses (e.g., 0.0, 0.025, 0.1 mg/kg/inf) from the first to second dose-response evaluations.

**Figure 7:**
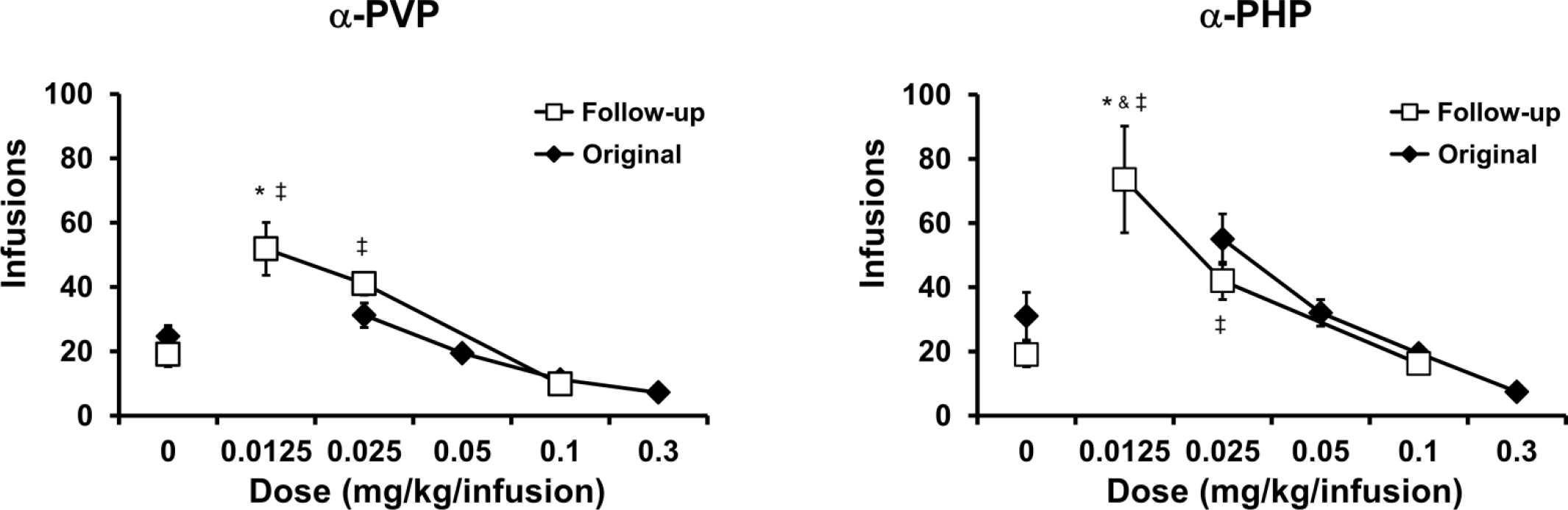
Mean (N=5; ±SEM) infusions of α-PVP and α-PHP obtained by female rats. A significant difference from vehicle is indicated with *, from the 0.025 mg/kg dose with & and from the 0.1 mg/kg dose with ‡.

## 4. Discussion

The study directly compared the psychostimulant effects of pentedrone, pentylone and methylone on measures of spontaneous locomotor activity and operant reward. Peak increases in activity rates were similar across all three compounds demonstrating similar efficacy; however, the locomotor effects of pentylone and methylone dissipated more quickly than those of pentedrone. Similar dose relationships were observed across male and female rat groups within and between the drugs, thus the outcome generalized across sex and across minor differences in age. The locomotor stimulant effects were of a similar magnitude to those observed for mephedrone, α-PVP and MDPV in prior investigations using similar methods, but appeared to last longer, i.e., more consistent with the duration of locomotor effects described for MDMA or methamphetamine (Aarde *et al*, 2013b; Miller *et al*, 2013a; Wright *et al*, 2012). Peak locomotor stimulant effects of α-PVP and MDPV were observed at 1.0 mg/kg, i.p. with reductions in activity observed after 5.6-10 mg/kg MDPV in a prior study (Aarde *et al*, 2013b). In this study pentedrone, pentylone and methylone exhibited a threshold for increased activity above 1 mg/kg and essentially linear increases in locomotor activity up to 10 mg/kg, i.p. suggesting less potency as locomotor stimulants compared with either α-PVP or MDPV. This was also the case for the induction of alternate behavior that interferes with locomotion which was not observed in this study at the highest doses of pentedrone, pentylone or methylone evaluated. In this, these three substituted cathinones appear to be more similar to mephedrone or MDMA and less similar to α-PVP or MDPV.

The intravenous self-administration (IVSA) study found a great deal of overlap in the dose-effect functions for pentylone and pentedrone, with a slight rightward shift of that for pentylone compared with pentedrone, indicating a potency difference. This contrasts with a prior MDPV/α-PVP comparison (Aarde *et al*, 2015a) where the 3,4-methylenedioxy motif appeared to convey no differences in potency or efficacy for the common structure. The function for methylone in this study reflected potency very similar to that of pentylone, however there was evidence for a reduction in peak responding (i.e., efficacy) of methylone compared with either pentylone or pentedrone. Evidence for the ascending and descending limbs of dose-effect functions was obtained for pentylone, pentedrone and methylone, thereby enhancing confidence that the full effective range was assessed. In contrast, only the descending limb for the α-PVP and α-PHP compounds was described over the 0.0125-0.3 mg/kg/infusion range which is consistent with higher potency as reinforcers compared with the other three compounds. The addition of the 0.0125 mg/kg dose in the follow-up study further confirmed this potency shift, but more importantly it confirmed that efficacy of α-PVP and α-PHP is as high as, or higher than, the other cathinones since peak responding was observed when a 0.0125 mg/kg/infusion dose was available. Although the primary focus here was not on α-PVP and α-PHP, which were included mostly as positive controls for the behavioral procedure, it is notable that α-PHP exhibited slightly more efficacy than did α-PVP. These compounds produce nearly identical inhibition of the DAT (Eshleman *et al*, 2017; Kolanos *et al*, 2015) and thus it would be of interest to determine what other pharmacological properties confer this difference in reinforcing efficacy. The reduced efficacy of methylone compared with pentedrone or pentylone is as would be predicted from the efficacy of methylone as a releaser of serotonin (Eshleman *et al*, 2013; Eshleman *et al*, 2017; Simmler *et al*, 2013; Simmler *et al*, 2014). Self-administration of methylone escalates under long (6 h) daily access conditions much more than does IVSA of MDMA despite only slightly enhanced IVSA under shorter (2 h) access conditions (Nguyen *et al*, 2016b; Vandewater *et al*, 2015). The dose-substitution data under short access conditions presented here suggest that pentylone and pentedrone would exhibit escalated intakes at least as great as those for methylone, and significantly higher than the IVSA of MDMA, under long access conditions. Nevertheless, the contrast of IVSA of methylone with MDMA under short- and long-access conditions cautions that any given pre-clinical model is limited in terms of predicting real-world abuse liability. Further differences or similarities across the entactogen cathinones might emerge with future studies that incorporate, e.g., choice procedures, IVSA under a Progressive Ratio schedule of reinforcement, etc.

Thermoregulatory impact was negligible for any of the three substituted cathinones under the tested ambient conditions and doses administered. This is consistent with a lack of temperature disruption in a prior study after 30 mg/kg, s.c., pentylone (Grecco *et al*, 2016); however, that report also found a pronounced hyperthermia after methylone (30 mg/kg, s.c.). It is unknown at present if cathinones categorically convey less risk of hyperthermia compared with amphetamine derivatives, but the 4-methyl substituted methcathinone (mephedrone) appears to mostly lower body temperature in conditions where MDMA produces hyperthermia (Miller *et al*, 2013a; Wright *et al*, 2012). Full determination of the risks of pentylone, pentedrone or methyone for unregulated temperature disruption would likely require higher doses administered at higher ambient temperatures.

In conclusion, pentedrone and pentylone are effective locomotor stimulants and function as reinforcers in the self-administration model. Potency differences between the two are subtle, showing that in this context the 3,4-methylenedioxy motif does not convey the substantial difference in behavioral effect that it does when added to methamphetamine. These compounds share the extended α-alkyl chain present on MDPV and α-PVP but appear to be less potent than either in comparison with prior findings. This suggests the pyrrolidine motif is critical to the high potency of those compounds compared with pentylone or pentedrone.

